# Acidic pH reduces *Vibrio cholerae* motility in mucus by weakening flagellar motor torque

**DOI:** 10.1101/871475

**Authors:** Nguyen T. Q. Nhu, Helen J. Wang, Yann S. Dufour

**Affiliations:** Department of Microbiology and Molecular Genetics, Michigan State University, East Lansing, Michigan, USA

**Keywords:** Vibrio cholerae, intestinal mucus, flagellar motility, motor torque, pH, microrheology

## Abstract

Intestinal mucus is the first line of defense against intestinal pathogens. It acts as a physical barrier between the epithelial tissues and luminal microbes. Enteropathogens, such as *Vibrio cholerae*, must compromise or circumvent the mucus barrier to establish a successful infection. We investigated how motile *V. cholerae* is able to penetrate mucus using single cell tracking in unprocessed porcine intestinal mucus. We found that changes in pH within the range of what has been measured in the human small intestine indirectly affect *V. cholerae* flagellar motor torque, and consequently, mucus penetration. Microrheological measurements indicate that the viscoelasticity of mucus does not change substantially within the physiological pH range and that commercially available mucins do not form gels when rehydrated. Finally, we found that besides the reduction in motor torque, El Tor and Classical biotypes have different responses to acidic pH. For El Tor, acidic pH promotes surface attachment that is mediated by activation of the mannose-sensitive haemagglutinin (MshA) pilus without a measurable change in the total cellular concentration of the secondary messenger cyclic dimeric guanosine monophosphate (c-di-GMP). Overall, our results support that the high torque of *V. cholerae* flagellar motor is critical for mucus penetration and that the pH gradient in the small intestine is likely an important factor in determining the preferred site of infection.

**Author summary:** The diarrheal disease cholera is still a burden for populations in developing countries with poor sanitation. To develop effective vaccines and prevention strategies against *Vibrio cholerae*, we must understand the initial steps of infection leading to the colonization of the small intestine. To infect the host and deliver the cholera toxin, *V. cholerae* has to penetrate the mucus layer protecting the intestinal tissues. However, *V. cholerae’s* interactions with intestinal mucus has not been extensively investigated. In this report, we demonstrate using single cell tracking that *V. cholerae* is able to penetrate native intestinal mucus using flagellar motility. In addition, we found that a strong motor torque is required for mucus penetration and, that torque is weakened in acidic environments even though the motor is powered by a sodium potential. This finding has important implications for understanding the dynamics of infection because pH varies significantly along the small intestine, between individuals, and between species. Blocking mucus penetration by interfering with *V. cholerae’s* flagellar motility, reinforcing the mucosa, controlling intestinal pH, or manipulating the intestinal microbiome, will offer new strategies to fight cholera.

## Introduction

*Vibrio cholerae* is the cause of an ongoing cholera pandemic with up to 4 million cases per year from regions of the world that do not have access to potable water [1]. Without proper rehydration and antibiotic treatments, the severe diarrhea triggered by the cholera toxin can be fatal [2]. Preventative measures and vaccines against *V. cholerae* have had partial success [3,4]. Therefore, cholera outbreaks are still a significant burden for populations living in developing regions or after natural disaster, such as Bangladesh and Haiti [1].

*V. cholerae* is represented by more than 200 serogroups that are endemic to sea and brackish waters and often found associated with copepods [5,6]. However, only the O1 and 0139 serogroups have been associated with cholera, the diarrheal disease in humans [7]. Within the O1 serogroup, the Classical biotype dominated the first 6 recorded cholera pandemics. The ongoing 7^th^ pandemic is dominated by the El Tor biotype, which has rapidly displaced Classical in the environment since its identification in 1905 [8,9]. Although similar, the two biotypes have differences in their genetic makeups, signaling dynamics, and behaviors [8,10–12]. The relative importance of their unique traits has not been fully elucidated yet.

*V. cholerae* colonize the mucus of the small intestine non-invasively. When reaching the intestinal crypts, *V. cholerae* secreted the cholera toxin, which targets epithelial cells to activate the chlorine channels proteins and consequently trigger a massive efflux of chlorine ions and water into the intestinal lumen. Many aspects of *V. cholerae* physiology and the regulation virulence factor expression have been investigated to recapitulate the dynamics of infection after ingestion, such as pili production [13], type 6 secretion system [14], quorum sensing [15], biofilm formation [16], and flagellar motility [17]. While these different behaviors have been shown to contribute to *V. cholerae* success during infection, the specific sequence of events and site-specific activities in the intestine are still under investigation.

Studies done on infant rabbits and mice indicate that in the early stage of infection planktonic *V. cholerae* cells are distributed throughout the small intestine. Then, the bacterial load drops in proximal and medial small intestine while the surviving cells preferentially colonize the distal small intestine [18,19]. Only, a small fraction of cells is able to penetrate the mucus layer protecting protect epithelial tissues. In the later stage of the infection, *V. cholerae* repopulates all parts of the small intestine [18,20]. It is likely that the dramatic change in the lumen conditions, which is triggered by the activity of the cholera toxin secreted by cells that reached the intestinal crypts first, enables *V. cholerae* to outgrow the host microbiota. It is still unclear what factors determine the preferential colonization of the distal small intestine.

Flagellar motility is essential for *V. cholerae* infection. Studies of transcription profiles and screens of mutant libraries during the infection of animal models and humans identified genes involved in chemotaxis and motility functions [21]. Non-motile *V. cholerae* mutants have reduced virulence and intestinal colonization [22–24]. In addition, previous work supports that protective immunity is mostly provided by mucosal antibodies that inhibit *V. cholerae* motility through bivalent binding the O-antigen [25]. However, motility may not be essential for survival and growth in the intestine since non-motile mutants do not appear to suffer a large competitive disadvantage when inoculated with motile *V. cholerae* [19].

The mucus layer protecting the intestinal tissue is difficult for bacteria to penetrate. Mucus is a complex hydrogel made of mucins (2-10% w/v), lipids, and DNA [26]. Mucins are large and highly glycosylated proteins cross-linked by disulfide bonds reinforced by hydrophobic interactions to form a tight mesh. The intestinal mucus layer is continuously renewed by secretion of highly O-glycosylated MUC2 mucin by goblet cells (240 ± 60 μm per hour) [27]. Consequently, mucus forms a selective diffusion barrier with constant flow rate, which can increase in response to threat such as the cholera toxin [28]. Histological analyses revealed that the inner part of the mucus layer is mostly free of bacteria [29]. In the small intestine, the mucus layer is thinner in the proximal part (~200 μm) than the distal part (~500 μm) [30]. These observations raise the questions of how *V. cholerae* is able to penetrate mucus and why it preferably infects the distal small intestine where the mucosa is thicker?

The pH gradient along the length of the small intestine may contribute to *V. cholerae*’s preferred site of infection. In humans, the proximal small intestine is slightly acidic (pH 6.3-6.5) while the distal part is slightly alkaline (pH 7.5-7.8) [31,32]. *V. cholerae* is able to grow between pH 6.5 and 9, but its preferred pH is that of sea water at ~8 [33]. *V. cholerae* also appears to be more motile at alkaline pH [34,35]. It has been reported that high gastrointestinal pH increases the susceptibility of *V. cholerae* infection [36] and that lactic acid producing bacteria, such as *Lactococcus lactis*, can help fight *V. cholerae* infections [37]. In contrast, acidic pH regulates the expression of virulence factors in *V. cholerae*. The production of cholerae toxin and toxin-coregulated pili is maximal at pH 6.6 [38,39]. The effects of intestinal pH on the dynamics of *V. cholerae* infection are not clearly established and have not been extensively investigated.

In this report, we investigated how pH affects mucus penetration by *V. cholerae*. To understand how *V. cholerae* interacts with mucus, we characterized both the swimming behavior of single *V. cholerae* cells in unprocessed intestinal mucus and the viscoelasticity of mucus at different pH. This analysis revealed that, although *V. cholerae* uses a sodium-driven flagellar motor, acidic pH weakens the motor torque. A weaker torque severely affects *V. cholerae*’s ability to penetrate intestinal mucus. We also show that the viscoelasticity of mucus does not change substantially within the physiological pH range. Therefore, the effect of pH on *V. cholerae* motility is likely an important factor in determining the preferred site of infection.

## Results

### pH affects the spread of *V. cholerae* colonies in soft agar

To test the effect of pH on *V. cholerae* motility, we measured the spread of colonies in M9 salts supplemented with pyruvate, tryptone, and 0.3% w/v agar (Figure 1A). The pH range was chosen based on what was measured in the human intestine where pH starts at 6 in the proximal part and gradually rises to 8 in the distal part [32]. Both the Classical and El Tor biotypes formed significantly smaller colonies at acid pH (Figure 1B). The colony morphology of the El Tor biotype was denser and rugged at the edge at all pH when compared to the Classical biotype. One of the differences between the two biotypes is that Classical does not elaborate the MshA (mannose-sensitive hemagglutinin) pilus that mediates cell attachment [40–42]. We inactivated *MshA* in the El Tor background to test if MshA affects colony morphology (Figure 1A). The colonies of the MshA mutant had smoother edges, spread more (Figure 1B), but remained dense like the wild type. Overall, *V. cholerae* spread faster in soft agar at alkaline pH. However, colony spreading is a function of cell motility and chemotaxis to self-generated chemical gradients but also growth rate [43–45]. In addition, *V. cholerae* growth is known to be sensitive to acidic pH [46].

**Figure 1.**
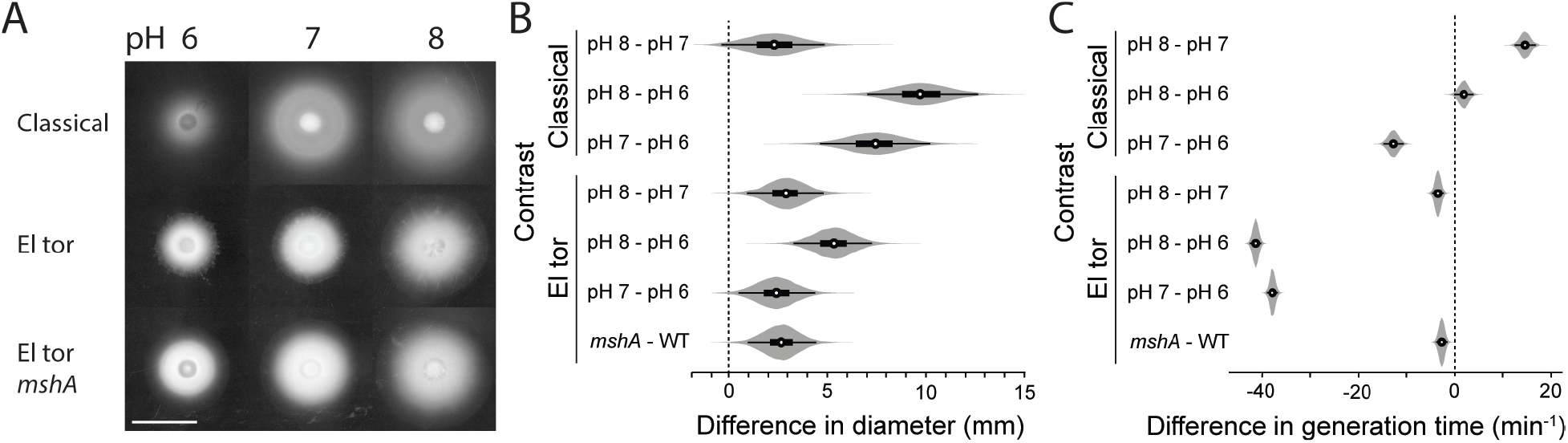
Effects of pH on the spreading of *V. cholerae* colonies in soft agar. (A) Representative colonies from the Classical and El Tor (wild type and *MshA*) biotypes at acidic, neutral, and alkaline pH (white bar is 10 mm). (B) Posterior probability distributions of the differences in colony size between different pH evaluated from a linear mixed-effect model (6 replicates for Classical and El Tor, 3 replicates for *MshA*). The model intercepts for the mean diameters are 21 mm for Classical and 15 mm for El Tor. (C) Posterior probability distributions of the differences in growth rate between different pH evaluated from a linear mixed-effect model (6 replicates). (circle: mean, thick line: 50% highest density interval, thin line: 95% highest density interval)

To test the effect of pH on cell growth in M9 salts supplemented with pyruvate, we measured the rate of increase in optical density from batch cultures at low cell density. At neutral pH, El Tor grew ~60% faster than Classical. pH had only a small effect on the generation time of Classical (Figure 1C). Classical grew faster at neutral pH (98 minutes generation time) and the generation times were very close between pH 6 (112 minutes) and 8 (114 minutes). El Tor grew fastest at pH 8 (59 minutes) but the difference was relatively small when compared to pH 7 (63 minutes). El Tor grew ~40% slower at pH 6 (103 minutes). The expression of MshA had a small but measurable effect on the generation time of El Tor. The effect of pH on generation time may explain why colony spreading was reduced for El Tor. However, these results do not explain why Classical was similarly affected by pH and spread faster than El Tor in soft agar. Therefore, we hypothesized that pH also affects *V. cholerae* flagellar motility.

### *V. cholerae* swims faster at alkaline pH

To test if *V. cholerae* change their motile behavior in different pH, we tracked single-cell swimming behavior in M9 salts between 2 glass coverslips (~10 μm in height). The diffusion coefficient of each trajectory was calculated to characterize the distribution of behaviors for each strain. To distinguish between motile and non-motile cells, we also tracked 1 μm polystyrene beads and a non-motile *flrA* mutant in the El Tor background. The diffusion coefficient of beads and non-motile cells was distributed between 0.1 and 10 μm^2^/s (Figure S1).

Therefore, trajectories with a diffusion coefficient above 10 μm^2^/s were categorized as motile cells. In Classical, most cells were highly motile near the end of the exponential growth phase. The average diffusion coefficient of the population increased upon transfer from the spent growth medium to fresh medium at all pH likely because of the replenishment of the energy source (addition of pyruvate to spent medium had an identical effect). However, Classical was most diffusive at alkaline pH (Figure 2A).

**Figure 2.**
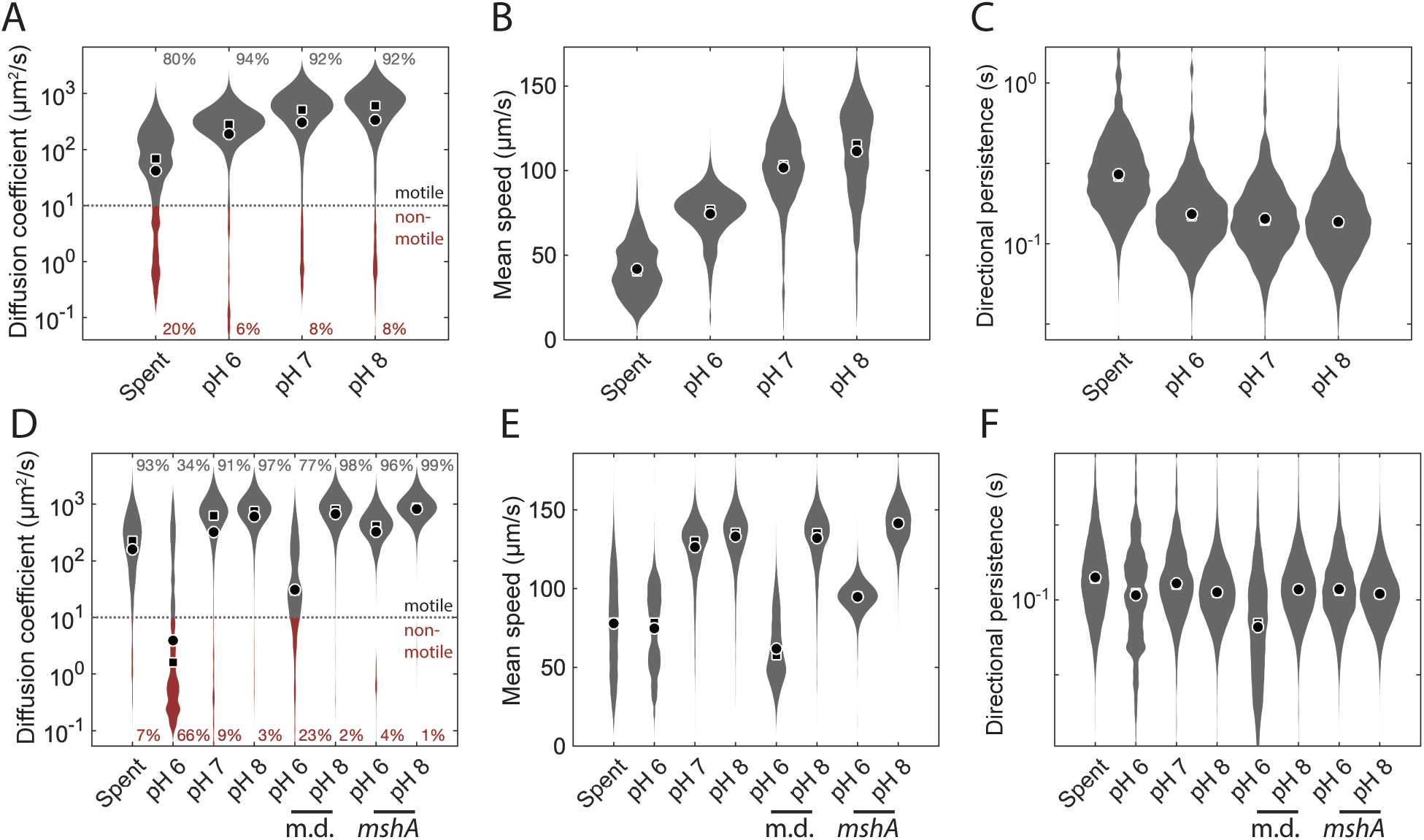
Effects of pH on *V. cholerae* flagellar motility. (A) Distributions of diffusion coefficients of Classical from single-cell trajectories in spent medium (Spent) or in fresh medium at different pH. Trajectory below 10 μm^2^/s were categorized as non-motile (red). (B) Distributions of mean swimming speed from the motile cell populations. (C) Distributions of trajectory directional persistence from the motile cell populations. (D) Distributions of diffusion coefficients of El Tor from single-cell trajectories in spent medium (Spent) or in fresh medium at different pH. Wild type El Tor was tracked in the presence of a mannose derivative (m.d.). A *MshA* mutant was also tracked (*MshA*). Trajectory below 10 μm^2^/s were categorized as non-motile (red). (E) Distributions of mean swimming speed from the motile cell populations. (F) Distributions of trajectory directional persistence from the motile cell populations. Each distribution represents 3 replicates combining between 2,000 and 10,000 individual trajectories (between 100 and 500 minutes of cumulative time). (circle: mean, square: median, standard error of the mean is too small to plot)

Both swimming speed and the frequency at which cell change direction by reversing the flagellar motor rotation affects the diffusion coefficient. However, analysis of the trajectories revealed that only swimming speed was affected by pH (Figure 2B). The average swimming speed changed from 74 μm/s to 111 μm/s between pH 6 and 8. On the other hand, the directional persistence of the cell trajectories did not change substantially, indicating that the reversal frequency of the flagellar motor was not affected by pH. Therefore, the reduction of swimming speed, caused by a weakening of the flagellate motor torque, is likely the main factor affecting the rate of colony expansion on soft agar as a function of pH for Classical (Figure 1).

Tracking of El Tor revealed a more complex behavioral response to change in pH. Upon transfer from the growth medium to pH 6, two third of the population became non-motile (Figure 2D) while at pH 7 and 8 the response was like that of Classical. The diffusion coefficients of non-motile cells were lower than what would be expected of cell subject to Brownian motion (Figure S1), indicating that cells were attaching to the glass surface. To confirm that cell attachment at pH 6 is mediated by MshA we also tracked El Tor in the presence of methyl α-D-mannopyranoside, a mannose derivative which blocks MshA interactions, and with a *mshA* mutant in the El Tor background. Addition of the mannose derivative only partially recovered motility while the *mshA* mutant is fully motile at pH 6 (Figure 2D). Therefore, El Tor activates MshA-mediated attachment at acidic pH but not at neutral or alkaline pH in our growth conditions. These results are consistent with the observation that the presence of MshA reduces the spread of colonies on soft agar (Figure 1).

In the absence of MshA, El Tor mean swimming speed was also severely reduced from 141 μm/s to 95 μm/s between pH 8 and 6 (Figure 2E). The directional persistence was unaffected indicating that pH does trigger a strong behavioral response (Figure 3F). These results are consistent with the response of Classical indicating that the mechanism underlying the reduction in swimming speed at acidic pH is the same in both biotypes. In addition, the reduction in speed is not the result of signaling activity through the chemotaxis pathway. c-di-GMP has been shown to regulate many behavioral responses in *V. cholerae*, including flagellar motility and surface attachment [47–49]. To test if the cytoplasmic c-di-GMP concentration changes after a shift in pH, we quantified the bulk c-di-GMP concentrations after transfer to buffer solution at different pH using mass spectrometry with El Tor sampled during the early stationary phase. No measurable change in the total c-di-GMP concentration within 1 μM could be attributed to change in pH (Figure S2). Our results cannot exclude that pH activates c-di-GMP signaling through localized pathways as previously demonstrated in *Escherichia coli* [50] or that c-di-GMP changed and returned to the pre-stimulus concentrations during the incubation period (15 minutes).

**Figure 3.**
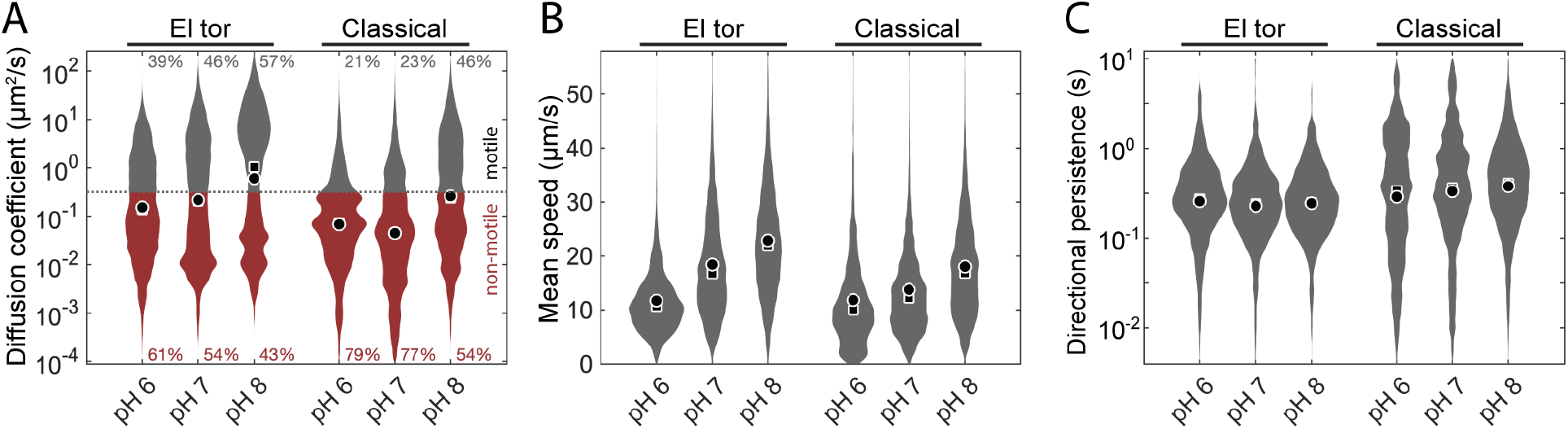
Effects of pH on motility through pig intestinal mucus. (A) Distributions of diffusion coefficients from single-cell trajectories in mucus buffered at different pH. Trajectory below 0.32 μm^2^/s were categorized as non-motile (red). (B) Distributions of mean swimming speed from the motile cell populations. (C) Distributions of trajectory directional persistence from the motile cell populations. Each distribution represents 8 to 12 replicates combining between 6,000 and 19,000 individual trajectories (between 1,000 and 2,600 minutes of cumulative time). (circle: mean, square: median, standard error of the mean too small to plot)

### *V. cholerae* penetrates mucus better at alkaline pH

To test if pH affects *V. cholerae*’s ability to penetrate mucus, we tracked fluorescently labeled cells in unprocessed mucus that was scraped from the medial part of a pig small intestine and equilibrated to different pH. As expected, the movement of both El Tor and Classical is severely impaired in mucus when compared to swimming in a liquid environment (Figure 3A, S1, and S3). A majority of cells were trapped in the mucus mesh. The rest of the population was able to swim through the mucus but would intermittently get trapped in the mucus mesh. Cells are not moving freely with a constant diffusion coefficient. Consequently, the reported diffusion coefficient represents an average over the trapped and free periods for each trajectory. The proportions of swimming cells and the average speeds increased in alkaline pH for both El Tor and Classical (Figure 3AB). The frequency at which cells change direction is not substantially affected by pH (Figure 3C).

Previous studies have characterized the behavior of *V. cholerae* in mucus reconstituted from commercially available purified mucin [51,52]. We repeated the characterization of single-cell motility in solutions of mucins from pig stomach or bovine sub-maxillary glands from Sigma-Aldrich. We used a 3% w/v concentration, which is comparable to native mucus [53][26]. We found that the bovine mucin solution quickly killed *V. cholerae* (survival rate from 0 to 0.06% after 45 minutes) unless dissolved in LB medium. We were unable to identify the source of toxicity. Ultimately, we found that the diffusion of *V. cholerae* was higher in the reconstituted pig and bovine mucin solutions than in our pig intestinal mucus samples (Figure S3). This result indicates that mucus reconstituted from purified mucins is different from native mucus and likely does not reconstitute a gel.

Overall, the tracking results are consistent with the effect of pH on swimming behavior in liquid. Therefore, the pH of the local intestinal environment is likely an important factor for the success of *V. cholerae* in colonizing the mucus layer in the small intestine of the human host. However, in addition to affecting the torque of the flagellar motor, alkaline pH may also affect the viscoelastic properties of mucus to facilitate cell translocation.

### pH has little effects on mucus viscoelasticity

To test if pH affects the structure of mucus, we tracked the motion of 1 and 0.2 μm fluorescent polystyrene beads coated with polyethylene glycol that were mixed in the same mucus samples used to track *V. cholerae*. The thermally driven diffusive behavior of beads is affected by the viscoelastic properties of mucus. Therefore, the loss (viscous) and storage (elastic) moduli of the mucus can be calculated from mean-squared displacement of the beads with respect to time using the generalized Strokes-Einstein relation [54]. This analysis indicates that the mucus viscoelasticity did not change substantially within the range of pH tested (Figure 4). The 1 μm beads had a sub-diffusive behavior (slope of the mean-squared displacement < 1) indicating that the large beads were trapped within the pores formed by the mucin mesh (Figure 4A) [26]. The 0.2 μm beads diffused more freely within the mucus (mean-squared displacement slope ~1) suggesting that the small beads are smaller than the average pore size of the mucin mesh that was previously estimated to be ~240 μm using electron microscopy (Figure 4C) [27,55]. Therefore, larger beads experience more viscoelastic stress (Figure 4BD). Consequently, the motion of 1 μm beads and similarly sized bacteria such as *V. cholerae* is severely diminished in mucus.

**Figure 4.**
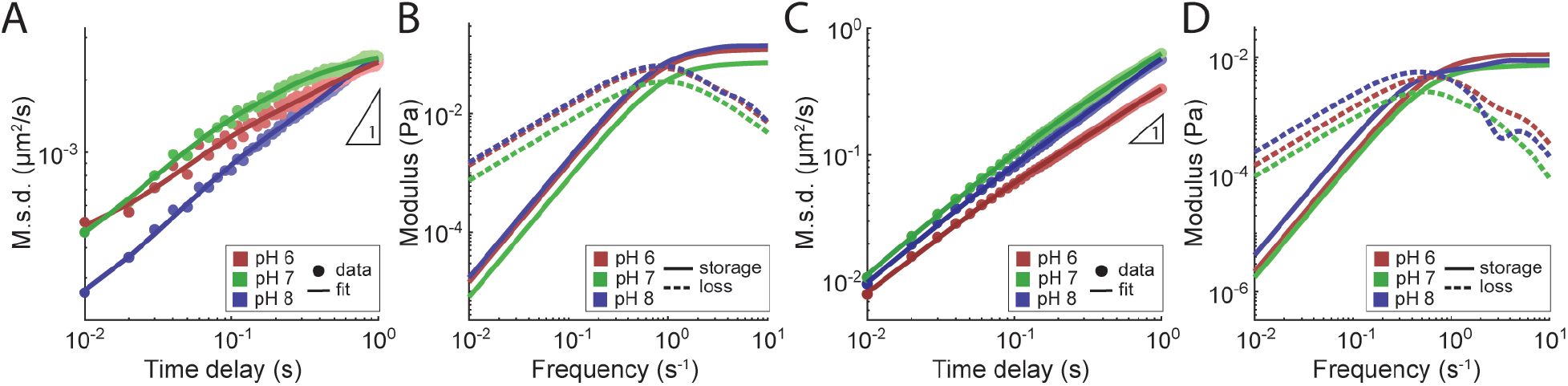
Passive microrheology of pig intestinal mucus. (A) Mean-squared displacement (M.s.d.) of PEG-coated 1 μm polystyrene beads with respect to time at different pH (colors). The data points (circles) are the average of trajectories from 4 to 6 replicates (10 to 25 individual trajectories). A polynomial fit the data (line) was used to calculate the storage and loss moduli using the generalized Strokes-Einstein relation. (B) Loss (dashed line) and storage (solid line) moduli of pig intestinal mucus at different pH (color) for 1 μm beads. (C) Mean-squared displacement (M.s.d.) of PEG-coated 0.2 μm polystyrene beads with respect to time at different pH (colors). The data points (circles) are the average of trajectories from 7 to 8 replicates (65 to 110 individual trajectories). (D) Loss (dashed line) and storage (solid line) moduli of pig intestinal mucus at different pH (color) from 0.2 μm beads.

We also characterized the viscoelasticity of the purified bovine and pig mucin solutions we used for characterizing the behavior of *V. cholerae*. Consistent with the behavior of swimming cells, the microbeads had purely diffusive trajectories indicating that that the solutions were viscous but not elastic (Figure S4A). The storage and loss moduli of the purified mucin solutions were lower than our pig mucus sample (Figure S4B). Therefore, the purified mucins failed to reconstitute the gel structure of native mucus. Incubation of non-motile *V. cholera* cells (El Tor *flrA*) with crude mucus for one hour did not produce a measurable change in the diffusion behaviors of cells and micro-beads (Fig S4CD).

Overall, we can conclude that the improvement seen in *V. cholerae* motility at alkaline pH is not the result of a change in the mucus viscoelasticity. Therefore, we propose that *V. cholerae*’s increased ability to penetrate the mucin mesh at alkaline pH is the result of higher torque from the flagellar motor.

### High motor torque is required for flagellar motility in hydrogels

*V. cholerae* uses a sodium-driven motor to rotate its flagellum. Therefore, pH is unlikely to have a direct effect on the motor torque and rotation speed. However, maintaining a strong sodium gradient across the cell membrane when the motor is rotating at high speed is energetically costly [56]. *V. cholerae* uses several sodium transporter but most of the sodium export is done by the NADH:quinone oxidoreductase (Na^+^-NQR) as part of the respiratory chain [57]. Activity of the Na^+^-NQR pump is strongest at alkaline pH while cells are respiring [58]. Therefore, we hypothesized that the reduction of swimming speed in spent medium or at acidic pH is the result of the reduction of the Na^+^-NQR pump activity.

To test this hypothesis, we measured the swimming speed of Classical in the presence of 2-n-Heptyl-4-hydroxyquinoline N-oxide (HQNO), a strong inhibitor of Na^+^-NQR activity [59]. The swimming speed of the motile cell population decreased in a dose-dependent manner with increasing concentration of HQNO (Figure 5A). The effect was more pronounced at acidic pH, indicating a possible synergistic interaction between pH and HQNO binding in the pump channel. However, swimming was not completely abolished even at 100 μM HQNO revealing the potential contribution of other sodium transporters. Overall, *V. cholerae* appears to struggle to maintain a strong sodium gradient at acidic pH, thus weakening the torque of the flagellar motor.

**Figure 5.**
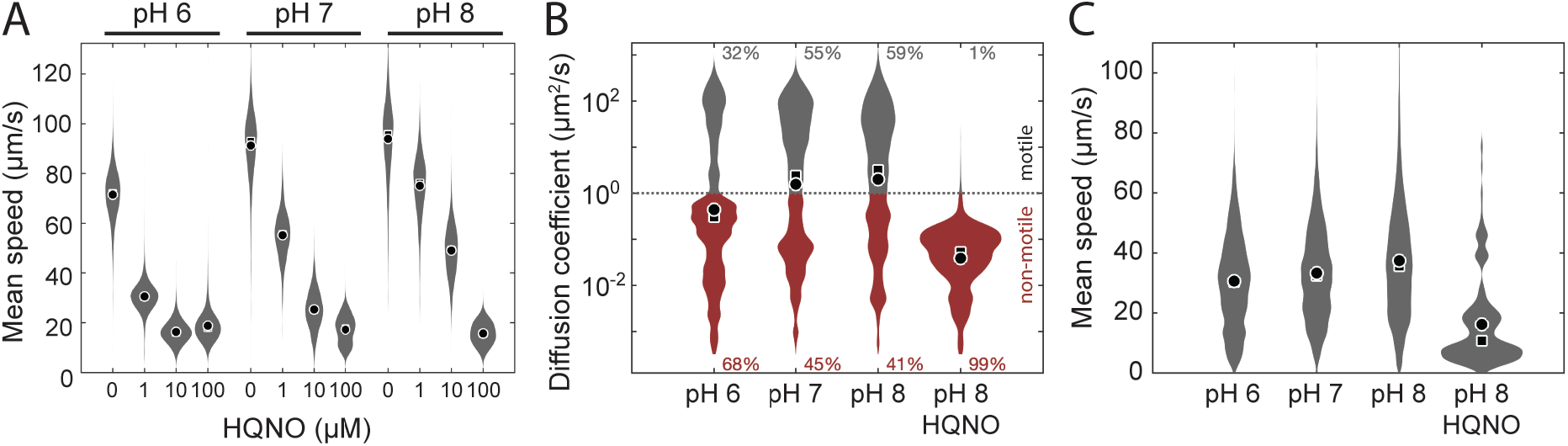
Effects of inhibiting the Na^+^-NQR pump activity on Classical motor torque. (A) Distributions of swimming speeds from the motile cell populations. Each distribution represents 3 to 4 replicates combining between 600 and 21,000 individual trajectories (between 150 and 450 minutes of cumulative time). (B) Distributions of diffusion coefficients from single-cell trajectories in agarose buffered at different pH and with the addition of 100 μM HQNO. Trajectory below 1 μm^2^/s were categorized as non-motile (red). (C) Distributions of mean swimming speed from the motile cell populations. Each distribution represents 6 to 12 replicates combining between 2,600 and 10,000 individual trajectories (between 380 and 600 minutes of cumulative time). (circle: mean, square: median, standard error of the mean too small to plot)

To test if reducing the flagellar motor torque would impact *V. cholerae*’s ability to swim through mucus, we tracked cells in the presence of HQNO. Unfortunately, mucus has a strong binding affinity to HQNO, which becomes unavailable to inhibit the Na^+^-NQR pump. Mucus has been previously shown to bind similar small molecules with high affinity [60]. Buffer containing 100 μM HQNO recovered after incubation with pig intestinal mucus also failed to inhibit *V. cholerae* swimming speed. Instead, we tested the effect of HQNO on *V. cholerae* motility through agarose gel. Low melting temperature agarose at 0.3% w/v forms a hydrogel similar to our pig intestinal mucus samples with slightly lower viscoelastic resistance (Figure S4). HQNO did not appear to interact with agarose and was able to inhibit the Na^+^-NQR pump. Consistent with mucus, the soft agar gel severely impaired but did not abolish *V. cholerae* motility and a higher proportion of cells were motile at alkaline pH (Figure 5BC). Addition of HQNO to agarose at pH 8 completely abolished motility of Classical supporting the hypothesis that *V. cholerae* requires a high motor torque to penetrate hydrogels.

## Discussion

In this work, we demonstrated that *V. cholerae* can penetrate intestinal mucus using its powerful flagellar motor. We extracted mucosa from a pig small intestine and characterized its viscoelastic properties to examine the physical challenge motile bacterial pathogens have to overcome to reach the epithelial tissues from the intestinal lumen. Crude intestinal mucus is a viscoelastic hydrogel with a pore size estimated to be between 200 nm and 1 μm from our microrheological analyses and previous imaging [27,55]. *V. cholerae* is small enough to swim through the mesh using flagellar motility. However, many cells were trapped in the mucin mesh and the effective diffusion coefficient of free-swimming cells was severely reduced when compared to swimming in liquid or purified mucin solutions that do not polymerize.

The diffusion of motile *V. cholerae* we observed in mucus may be sufficient to reach epithelial tissues during infection of the human small intestine. Previous studies have indicated that directional motion controlled by chemotaxis is not required for *V. cholerae* to infect the host [19,24,61]. Therefore, *V. cholerae* is likely performing a diffusive random walk through the mucosa. The typical thickness of mucus in the human small intestine is in the order of a few hundred μm and grows about 240 μm per hour [27]. The typical first-passage time of a diffusive trajectory can be calculated as the square of the distance to cross divided by twice the diffusion coefficient [62]. From our results, we estimate that the typical time *V. cholerae* would take to penetrate 400 μm of the small intestine mucosa at pH 8 is about 2 hours, which is very close to the time it takes to grow mucosa of that thickness. Therefore, in the absence of factors that interfere with flagellar motility, *V. cholerae* is intrinsically capable of overcoming the physical barrier formed by intestinal mucus using flagellar motility only.

In the conditions we tested, incubation of *V. cholerae* in crude pig mucus did not produce measurable changes in the mucus bulk physical properties. A previous study proposed that *V. cholerae* shears or loses its flagellum in the presence of bovine mucin and initiates the expression of virulence factors [51]. In this study, we found that *V. cholerae* rapidly dies in bovine mucin solutions unless dissolved in rich media (likely quenching an unidentified toxic compound). Dead cells showed the expected Brownian motion consistent with previous observations [51]. Solutions of purified pig gastric mucins did not show toxicity. We found that *V. cholerae* can grow in crude pig intestinal mucus and that the motile behavior is steady. These results indicate that, beside the physical interactions with the mucus mesh, there were no measurable biological interactions between *V. cholerae* and mucus in our experimental conditions.

*V. cholerae*’s flagellar motor torque is an important factor for mucus penetration. More cells were trapped in the mucin mesh and the diffusion coefficient of swimming cells decreased when motor torque was reduced. Because *V. cholerae*’s flagellar motor is powered by the transmembrane sodium gradient, the effect of environmental pH on motor torque is indirect. It has been shown that *V. cholerae*’s main sodium pump, Na^+^-NQR, has reduced activity at acidic pH [58]. Inhibiting Na^+^-NQR with HQNO had the same effect as acidic pH on *V. cholerae*’s motility in hydrogels. In addition, a previous study reported that mutant strain lacking NqrA (a subunit of the Na^+^-NQR complex) is defective at colonizing infant mice [63]. Therefore, our current model is that *V. cholerae* has difficulty maintaining a strong sodium motive force at acidic pH, reducing the cells capacity to penetrate mucus and reach the epithelium. In the human small intestine, pH gradually increases from the proximal to the distal parts, becoming alkaline in the ileum [32]. Therefore, the pH gradient along the small intestine likely plays an important role in determining the preferred site for mucus colonization by *V. cholerae*.

The dynamics of infection of the human small intestine by *V. cholerae* has not been firmly established, partially because of the limitations of existing animal models [64]. Our results identified additional factors to consider when interpreting experimental results. The small intestine of suckling mice and infant rabbits are likely to have different pH profiles and thinner mucosal tissues from humans, but limited information is currently available. The early infection steps may differ significantly between animal models and humans. Previous studies provided conflicting evidence supporting the role of flagellar motility during infection [65]. Some studies found that non-motile cells are less infectious [25,66], while others reported that there is no difference and that non-motile cell can reach the epithelial crypts in infant mice [19]. Therefore, the route to the epithelium may vary between experimental models. It is also possible that when animals are inoculated with libraries of mutants, non-motile mutants are benefit from the activity of motile cells that can penetrate the mucosa and deliver the toxin. The lumen conditions change drastically after the cholera toxin is delivered. The rapid flux of ions and water in the lumen disrupts the mucosa, mixes the lumen, and create favorable conditions for *V. cholerae*.

In humans, acquired immunity to *V. cholerae* appears to target flagellar motility [25,67] and heterozygous carriers of cystic fibrosis are more resistant to *V. cholerae* infection [68]. In addition, acid producing commensal bacteria provide protection against *V. cholerae* [37,69,70], which may affect *V. cholerae*’s ability to penetrate mucus rather than survival. These observations are consistent with an infection model where flagellar motility is critical at the very early stage of infection. Successful penetration of the mucosa has a low chance of success, but the disruption triggered by the delivery of the cholera toxin to the epithelium gives a growth advantage to *V. cholerae* over the rest of the gut microbiota. Further characterization of factors that affect *V. cholerae*’s flagellar motility through mucus are necessary to support this model and determine if therapeutic strategies to prevent mucus penetration would be effective. Improving or complementing the mucus as a physical barrier to block the access of pathogenic bacteria to epithelial tissues may have several advantages over the use of antibiotics. This approach would apply less selective pressure to evolve or acquire antibiotic resistance and it would not disrupt the commensal microbiota.

## Methods

### Bacterial strains

*V. cholerae* strains used in this study were El Tor C6706str2 [71] and Classical O395 [72] biotypes. Our wild type El Tor strain has a functional *lux*O gene. Strains were fluorescently labelled with the expression of the green fluorescent protein expressed from a constitutive cytochrome c *V. cholerae* promoter on a p15a plasmid derivative (gift from Dr. Chistopher Waters). The inactivation of *mshA* in the El Tor background was generated by recombining genomic DNA of mutant EC4926 from the defined transposon mutant library [73] using natural transformation [74]. The El Tor *flrA* mutant was generated from previous work [75].

### Growth conditions

M9 minimal salts (52 mM Na_2_HPO_4_, 18 mM K_2_HPO_4_, 18.69 mM NH_4_Cl, 2 mM MgSO_4_) were supplemented with 10 μM FeSO_4_, 20 μM C_6_H_9_Na_3_O_9_ and 36.4 mM Sodium pyruvate. The pH of the growth medium was adjusted with HCl to the desired value. *V. cholerae* was grown shaking (200 r.p.m.) in liquid cultures at 37°C. Kanamycin was added to 50 μg/ml when needed. For all experiments, *V. cholerae* cultures were sampled at early stationary phase (1.9 × 10^9^ c.f.u./mL). Soft agar plates were prepared with the same medium with the addition of 0.1% w/v Tryptone and 0.3% w/v Bacto agar (BD). Plates were inoculated with 5 μl of saturated liquid cultures (5.8 × 10^6^ cells) on the agar surface and incubated at 37°C for 12 hours before measuring colony size.

### Colony size and growth rate analyses

The posterior probability distributions for the treatment effects on colony size from soft agar plate were calculated using Bayesian sampling of a linear mixed-effect model with Gaussian distributions taking into account, pH, parent strain, genotype, and variability between replicates (Colony diameter ~ 1|Strain:pH + 1|Replicate + 1|Strain:msha) using the RSTAN [76] and BRMS packages [77,78] in R [79]. The plots were generated using the ggplot2 [80] and tidybayes [81] packages. The growth rates of bacterial cultures were calculated by recording the change in optical density at 590 nm of 200 μL cultures in 96-well plates (Corning, CLS3595) using a Sunrise plate reader (Tecan Trading AG, Switzerland). Cultures were inoculated with 1.6 × 10^6^ c.f.u./mL cells in the exponential growth phase and incubated at 37°C with intermittently shaking every 10 mins for 24 hours. Precautions were taken to limit evaporation. The generation time was calculated for each growth curve and the posterior probability distribution for each treatment across replicates was estimated using the same model (Generation time ~ 1|Strain:pH + 1|Replicate + 1|Strain:msha).

### c-di-GMP quantification

The concentration of c-di-GMP was measured as previously described [82]. Briefly, 2×10^8^ cells sampled during the exponential growth phase were collected on a PTFE membrane filter (0.2 μm) from each condition. Membranes were submerged and mixed in extraction buffer (40% v/v acetonitrile, 40% v/v methanol, 0.1 N formic acid) for 30 minutes. The extraction solution was spiked with a known amount of N^15^-labeled c-di-GMP to normalize sample loss across samples during extraction. Non-soluble cell debris were separated by centrifugation (21,130 r.c.f. for 2 minutes). The soluble fractions were dried in vacuum overnight and resuspended in 100μl distilled water prior to identification and quantification using mass spectrometry (Quattro Premier XE mass spectrometer, Waters Corp.). c-di-GMP and N^15^-labeled c-di-GMP were detected simultaneously at *m/z* 699.16 and at *m/z* 689.16 respectively. The posterior probability distributions for the treatment effects were calculated using Bayesian sampling of a linear mixed-effect model with Gaussian distributions ([c-di-GMP] ~ 1|pH + 1|Replicate) using the RSTAN [76] and BRMS packages [77,78] in R [79]. The plots were generated using the ggplot2 [80] and tidybayes [81] packages.

### Single-cell tracking

*V. cholerae* cells were tracked in liquid medium following the protocol previously described [83]. Briefly, *V. cholerae* cells in the early stationary growth phase were diluted to 1.9 × 10^7^ cells/mL in fresh medium adjusted to pH 6, 7, or 8. Methyl α-D-mannopyranoside was added to 100 mM when indicated. Cells were incubated shaking at 37°C for 15 minutes before tracking to allow for adaptation of the chemotaxis response. Polyvinylpyrrolidone (PVP) was added at 0.05% w/v to the samples to prevent attachment on the glass slide. 6 μl of each sample dropped on a glass slide and trapped under a 22 × 22 mm, #1.5 coverslip sealed with wax and paraffin to create a thin water film (10±2 μm) for video microscopy. For tracking in mucus or low-melting temperature agarose, a 130 μm spacer was added between the slide and the coverslip and fluorescently labelled cells were used. The samples were kept at 37°C during tracking. Images of swimming cells were recorded using a sCMOS camera (Andor Zyla 4.2, Oxford Instruments) at 20 frames per second using a 40X objective (Plan Fluor 40x, Nikon Instruments, Inc.) mounted on an inverted microscope (Eclipse Ti-E, Nikon Instruments, Inc.). Cell were illuminated using phase contrast in liquid or epifluorescence in mucus and agarose. Images were analyzed to detect and localize cells using custom scripts [83] and cell trajectories were reconstructed using the μ-track package [84]. The analysis and plots of the cell trajectory statistics were done in MATLAB (The Mathworks, Inc.) as previously described [83].

### Passive microrheology of mucus and agarose gel

The viscoelasticity of mucus and agarose were measured by tracking the passive diffusion of 1 μm and 200 nm fluorescent polystyrene beads (F8814 and F8810, ThermoFisher Scientific). To prevent electrostatic or hydrophobic interactions between the beads and the gels, beads were coated with polyethylene glycol (PEG MW 2,000Da). Coating was done by crosslinking carboxyl groups on the surface of the beads with diamine-PEG following the previously described protocol [85]. Beads 0.5% w/v and Triton (X-100, Sigma-Aldrich) 0.01% w/v were added to samples and mixed gently. Epifluorescence signal from the beads were recorded using a sCMOS camera (Andor Zyla 4.2, Oxford Instruments) at 100 frames per second using a 100X objective (Plan Fluor 100x, Nikon Instruments, Inc.) and a 1.5X multiplier mounted on an inverted microscope (Eclipse Ti-E, Nikon Instruments, Inc.). Images were analyzed to detect and localize beads using custom scripts and trajectories were reconstructed using the μ-track package [84]. The beads trajectories were manually inspected to remove artifact and erroneous linking. Systematic drift of the trajectories was corrected prior to calculating the bead average mean-squared displacement (MSD) and velocity autocorrelation (VAC) as a function of time. The VAC was fitted to a degree six polynomial multiplied by an exponential decay function. The VAC function was integrated according the Green-Kubo relation [86,87] to obtain a function that can also be fitted to the MSD with the same parameters. The VAC and MSD were fitted simultaneously using nonlinear least-square regression to separate the dynamic properties of the beads from the tracking noise. The fitted parameters were then used to calculate the storage (G’) and loss (G”) moduli of the sample according to the generalized Stokes–Einstein equation [88]. The analysis and plots of the bead diffusive behavior were done in MATLAB (The Mathworks, Inc.).

### Mucus preparation

Small intestines were obtained from a freshly slaughtered adult pig at the Meat Lab at Michigan State University (USDA permit #137 from establishment #10053). The animal was slaughtered as part of the normal work of the abattoir according to the rules set by the Michigan State University Institutional Animal Care and Use Committee (IACUC). The small intestines were acquired from the abattoir with prior consent. The mucosa was gently scraped from the medial part of small intestine and frozen in liquid nitrogen before storage at −80°C. For each experiment, mucus samples were warmed to 37°C temperature and equilibrated for 1 hour in 10 volumes excess of M9 salts buffered to the desired pH. Bovine submaxillary gland mucin (M3895, Sigma-Aldrich) solutions was prepared at 3% w/v in LB medium adjusted to pH 8.0 with sodium hydroxide. Non soluble particles were separated from the preparation by centrifugation at 21,130 r.c.f for 10 minutes. Porcine stomach (M2378, Sigma-Aldrich) mucin solution was prepared at 3% w/v in M9 salts at pH 8.0. The survival rate of *V. cholerae* in BSG was calculated by enumerating colonies on LB agar plate supplemented with 50 μg/ml kanamycin. Fluorescent beads were added to the samples at 0.15% w/v and gently mixed.

## Acknowledgements

This work was supported by Michigan State University funds and the National Science Foundation under Grant No. 1714612 to Y.S.D. We thank C. M. Waters and V. J. DiRita for gifting *V. cholerae* strains, the Mass Spectrometry Core at MSU, J. L. Franklin for assistance with data analysis, J. S. Lee for assistance with c-di-GMP extraction, and B. Hsueh for mass spectrometry trace analysis.

## Author contributions

Y. S. D and N. T. Q. N. conceived and designed the study. N. T. Q. N. and H. J. W. performed the experiments. Y. S. D. and N. T. Q. N. analyzed the data and wrote the manuscript with input from all authors.

## Supplementary materials

Video S1: Fluorescent *V. cholerae* El Tor motility in crude pig intestinal mucus.

---

**Figure S1.**
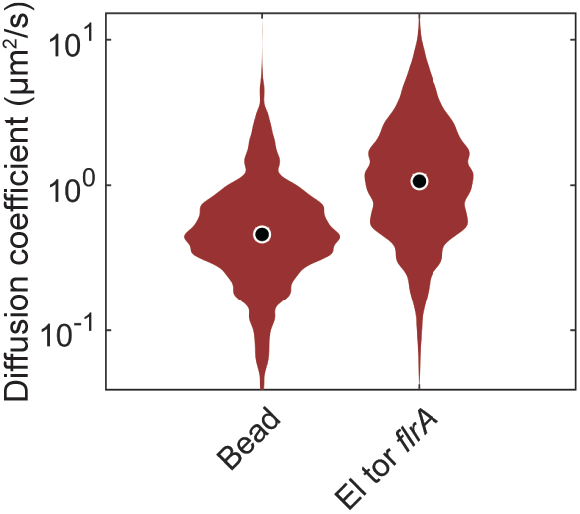
Distributions of diffusion coefficients of 1 μm beads and non-motile *V. cholerae* El Tor cells (*flrA*) in buffer solution.

**Figure S2.**
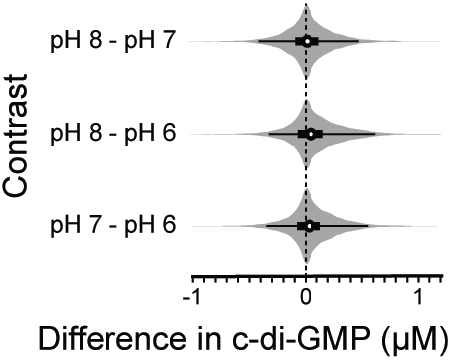
Change in c-di-GMP concentrations in response to change in pH in *V. cholerae* El Tor. Posterior probability distributions of the differences in c-di-GMP concentrations evaluated from a linear mixed-effect model. The model intercepts for the mean c-di-GMP concentrations is 3.90 μM. The analysis was done with 6 replicates. (circle: mean, thick line: 50% highest density interval, thin line: 95% highest density interval)

**Figure S3.**
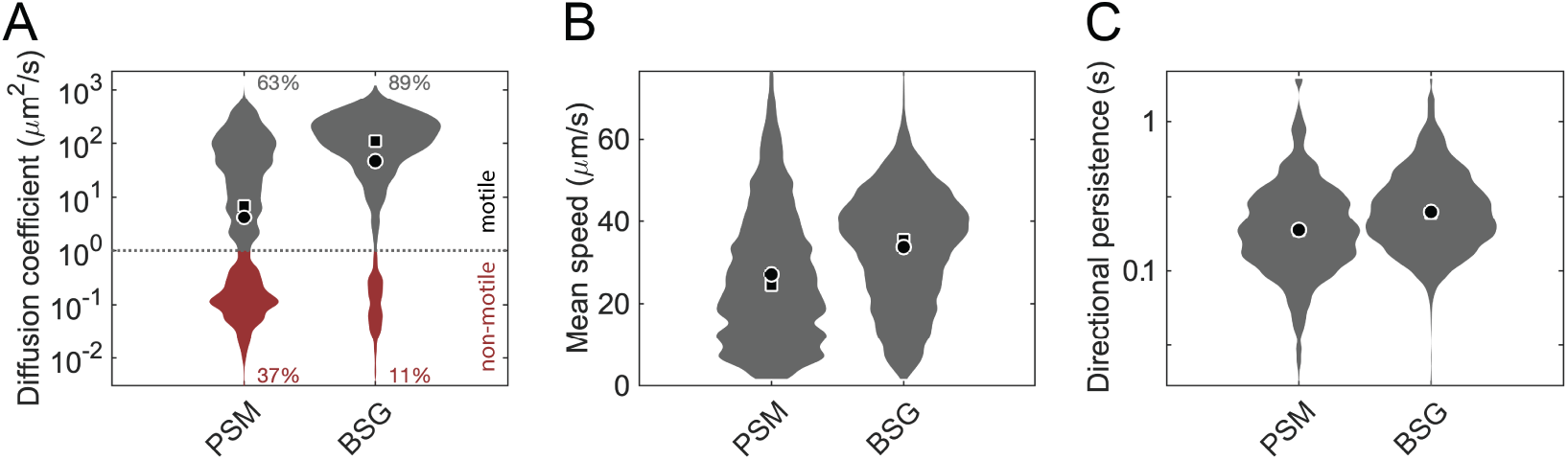
*V. cholerae* El Tor motility through commercial purified mucins. (A) Distributions of diffusion coefficients from single-cell trajectories in 3% w/v mucin from porcine stomach Type II in M9 salts (PSM) and 3% w/v mucin from bovine submaxillary glands Type I-S in LB (BSG) buffered at pH 8. Trajectory below 1 μm^2^/s were categorized as non-motile (red). (B) Distributions of mean swimming speed from the motile cell populations. (C) Distributions of trajectory directional persistence from the motile cell populations. Each distribution represents 4 replicates combining between 5,000 to 8,000 individual trajectories (between 124 and 170 minutes of cumulative time). (circle: mean, square: median, standard error of the mean too small to plot)

**Figure S4.**
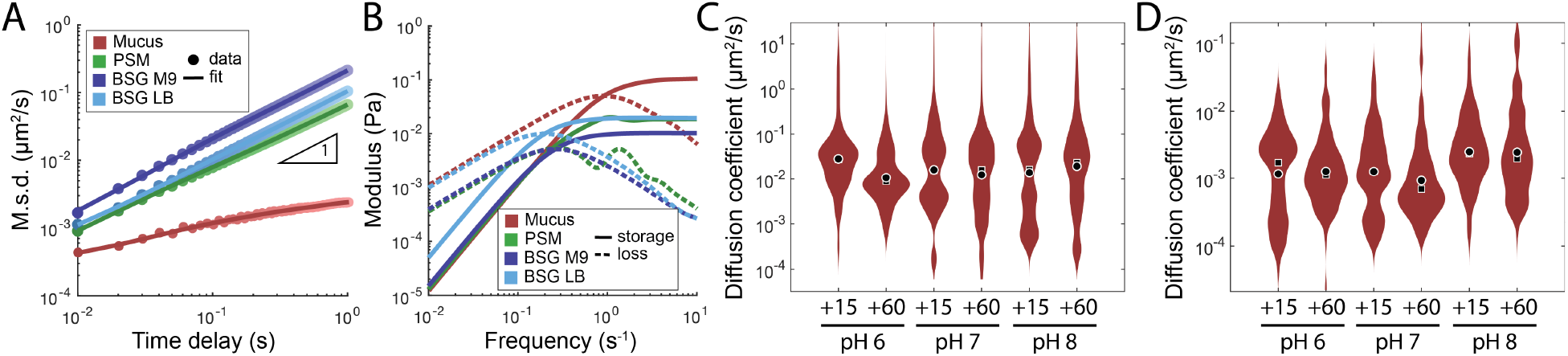
Passive microrheology of BSG and PSM and effects of V. cholera on crude mucus. (A) Mean-squared displacement (M.s.d.) of PEG-coated 1 μm polystyrene beads with respect to time in different mucus preparations (colors). The data points (circles) are the average of trajectories from 3 to 6 replicates (12 to 25 individual trajectories). A polynomial fit the data (line) was used to calculate the storage and loss moduli using the generalized Strokes-Einstein relation. (B) Loss (dashed line) and storage (solid line) moduli of different intestinal mucus preparations (color) for 1 μm beads. (C) Diffusion coefficient distributions of *V. cholerae* El Tor *flrA* in crude mucus after 15- and 60-minutes incubation at different pH. (D) Diffusion coefficient distributions of 1 μm polystyrene beads in the same mucus samples as (C). Each distribution represents one biological replicate with 600 to 900 individual trajectories (150 to 225 minutes of cumulative time). (circle: mean, square: median)

**Figure S5.**
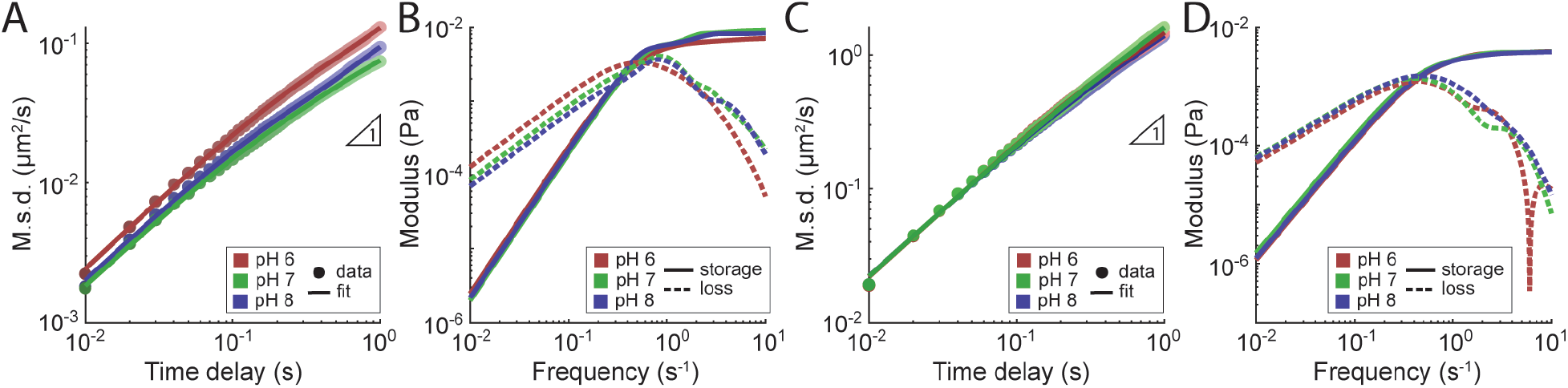
Passive microrheology of 0.3% w/v agarose. (A) Mean-squared displacement (M.s.d.) of PEG-coated 1 μm polystyrene beads with respect to time at different pH (colors). The data points (circles) are the average of trajectories from 7 to 8 replicates (28 to 29 individual trajectories). A polynomial fit the data (line) was used to calculate the storage and loss moduli using the generalized Strokes-Einstein relation. (B) Loss (dashed line) and storage (solid line) moduli of agarose gel at different pH (color) for 1 μm beads. (C) Mean-squared displacement (M.s.d.) of PEG-coated 0.2 μm polystyrene beads with respect to time at different pH (colors). The data points (circles) are the average of trajectories from 6 to 8 replicates (88 to 93 individual trajectories). (D) Loss (dashed line) and storage (solid line) moduli of agarose gel at different pH (color) from 0.2 μm beads.

